# Widespread temporal niche partitioning in an adaptive radiation of cichlid fishes

**DOI:** 10.1101/2024.05.29.596472

**Authors:** Annika L. A. Nichols, Maxwell E. R. Shafer, Adrian Indermaur, Attila Rüegg, Rita Gonzalez-Dominguez, Milan Malinsky, Carolin Sommer-Trembo, Laura Fritschi, Walter Salzburger, Alexander F. Schier

## Abstract

The partitioning of ecological niches is a fundamental component of species diversification in adaptive radiations. However, it is presently unknown if and how such bursts of organismal diversity are influenced by temporal niche partitioning, wherein species avoid competition by being active during different time windows. Here, we address this question through profiling temporal activity patterns in the exceptionally diverse fauna of cichlid fishes from African Lake Tanganyika. By integrating week-long longitudinal behavioural recordings of over 500 individuals from 60 species with eco-morphological and genomic information, we provide two lines of evidence that temporal niche partitioning facilitated this massive adaptive radiation. First, Tanganyikan cichlids exhibit all known circadian temporal activity patterns (diurnal, nocturnal, crepuscular, and cathemeral) and display substantial inter-specific variation in daily amounts of locomotion. Second, many species with similar habitat and diet niches occupy distinct temporal niches. Moreover, our results suggest that shifts between diurnal and nocturnal activity patterns are facilitated by a crepuscular intermediate state. In addition, genome-wide association studies indicate that the genetics underlying activity patterns is complex, with different clades associated with different combinations of variants. The identified variants were not associated with core circadian clock genes but with genes implicated in synapse function. These observations indicate that temporal niche partitioning contributed to adaptive radiation in cichlids and that many genes are associated with the diversity and evolution of temporal activity patterns.

## INTRODUCTION

Adaptive radiations are characterised by rapid species diversification as a consequence of niche specialisation^1–3^. For example, the beaks of Darwin’s Finches are highly specialised for different diets^3^, the varied limbs of anole lizards allow access to different sections of their tropical forest habitat^4^, and the diverse body and jaw shapes of African cichlid fishes match their habitats and diets^5^. Ecological niches can also be of temporal nature^6^. For example, species can specialise to be more active during certain time windows, such as the day, the night, or twilight periods. While such cases of temporal niche partitioning have been well documented between distantly-related species that coexist in sympatry^7–9^, it is largely unknown whether variation in circadian temporal activity patterns can contribute to niche specialisation in the course of adaptive radiations. Additionally, little is known about the molecular or genetic mechanisms that underlie different circadian activity patterns.

Cichlid fishes (Cichlidae) are one of the most species-rich families of vertebrates^10^. They commonly diversify through adaptive radiation, as seen across their geographic distribution and especially in the East African Great Lakes^5,11,12^. At approximately 9-12 million years of age Lake Tanganyika is the oldest of the East African Great Lakes. This lake harbours the most diverse adaptive radiation of cichlid fishes, displaying huge disparity in morphology, ecology, behaviour and genetics^12^. Anecdotal observations have suggested that cichlid species are mostly diurnal, and rely heavily on visual systems and colouration patterns associated with daylight^13^. However, there are reports that some species of cichlids are nocturnal^14^, or harbour traits often associated with a nocturnal lifestyle such as large eyes and an expanded lateral line^13,15,16^. These observations raise the possibility that cichlid radiations have been facilitated by temporal niche partitioning, with eco-morphologically similar species occupying distinct temporal activity patterns. Here we test this hypothesis by examining temporal activity patterns and their genetic underpinnings in the cichlid fishes of Lake Tanganyika.

## RESULTS

### Activity patterns are highly diverse in Tanganyikan cichlids

To investigate the temporal activity patterns in the cichlid fish fauna of Lake Tanganyika, we developed a behavioural tracking paradigm to follow hundreds of individually-housed fish over week-long periods in a reductionist and controlled laboratory setting, or “common garden”^17–20^. We collected data for 60 species from the adaptive radiation of cichlid fishes in Lake Tanganyika, including representatives from 9 out of the 12 tribes, and covering all trophic levels and all major diet guilds (**Fig. 1a**)^5^. We examined both the largest and smallest cichlids in the world (*Boulengerochromis microlepis* [abbreviated as Boumic] and *Neolamprologus multifasciatus* [Neomul], see **Supplementary Data 1** for list of species names and abbreviations), and species from all major biotypes in that lake, resulting in a phylogenetically, ecologically and morphologically diverse and representative set of species. In our experiments, we continuously tracked each individual fish for 6 days and 6 nights at high temporal resolution (10hz) within tanks separated by mesh dividers (**Fig. 1b**). This setup allowed fish to interact with each other through visual and olfactory cues, while facilitating robust tracking of individuals. Up to 14 adult individuals per species were tracked (average 9 individuals, with a range from 2-14 see **Supplementary Data 1**). Online tracking by custom python code recorded the position and speed of each fish.

**Figure 1.**
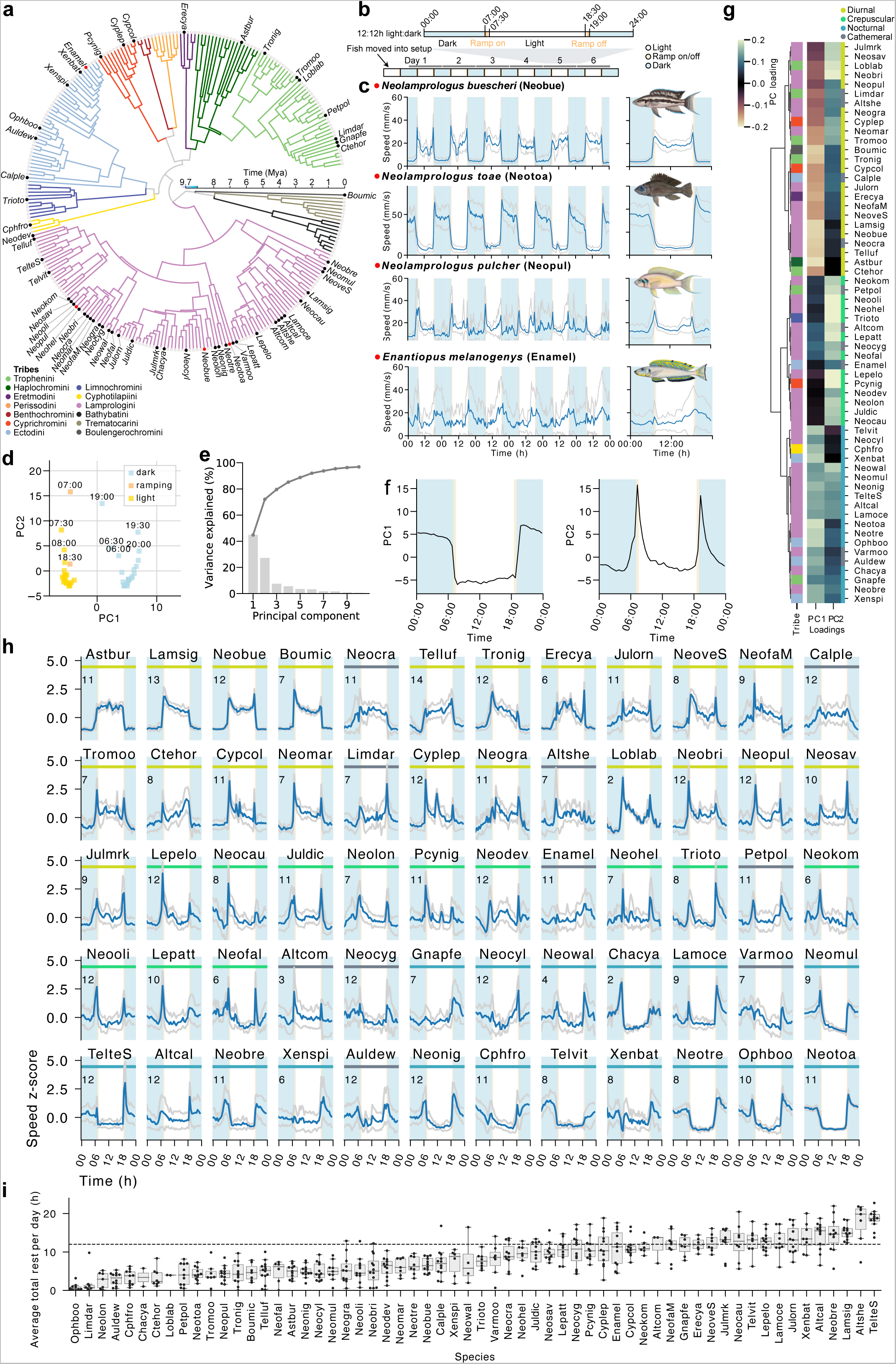
Cichlids occupy all known temporal activity niches and display extensive variation in total rest across species. **a**, Time calibrated phylogenetic tree of cichlid species from the Lake Tanganyikan radiation with branches coloured according to tribe. Labelled species were included in our behavioural screen. Red dots indicate example species shown in panel (c). **b**, Schematic of the timeline and light cycle for the behavioural assays. **c**, The weekly and daily average speed traces (mean +/- SD) of four example species. **d**, PCA analysis of the daily speed averages across the 60 species separates out the 30 min time bins by light state. **e**, The variance explained by the first 10 principal components. The overlaid line shows the cumulative sum of variance explained. **f**, Plot of PC1 and PC2 values. **g**, The clustered loadings of PC1 and PC2 are plotted along with the tribes (same colour key as in (a)), and diel guilds. **h,** The z-score normalised average daily speed traces for each of the 60 cichlid species assayed ordered by PC1 loadings. Coloured bar indicates the temporal activity pattern of the species (same colour key as in (g)). Numbers of animals assayed for each species are shown in the top left corner in each plot. **i**, Average daily total rest for each species, each dot shows the average for one individual. Species names are abbreviated using a six-letter code following Ronco *et al.* 2021 (**Supplementary Data 1**).

We uncovered a remarkable diversity of activity patterns across these closely related species (**Fig. 1** and **Extended Data Fig. 1, Supplementary Data 2).** Patterns ranged from diurnal (e.g. in *N. buescheri* [Neobue]) to nocturnal activity (e.g. in *N. toae* [Neotoa]) to peaks in activity at dawn and dusk (e.g. in *N. pulcher* [Neopul]) to no strong preference in daily activity (e.g. in *Enantiopus melanogenys* [Enamel]) (**Fig. 1c**). Our results are largely consistent with anecdotal observations from the wild available for some of the species. For example, *N. toae* is known to feed on insect larvae during the night, has very large eyes relative to its body size, and an expanded lateral line system^13^. In comparison, its congener *N. buscheri* has a smaller relative eye size than *N. toae*, is colourful and known to be aggressive and active during the day^13^. These examples suggest that our measured lab-based activity patterns can match known physiological adaptations indicative of natural temporal activity patterns.

To determine the major axes of variation in behavioural activity patterns across species, we performed dimensionality reduction using Principal Component Analysis (PCA) of their 30min binned average daily swimming speed, with species as features. This approach revealed two components that explained 72% of the variation of the daily activity patterns (PC1 explained 45% and PC2 explained 27% of variance, while PC3 only explained 8%) (**Fig. 1d-e**). The PC1 axis separated day timepoints (when lights were on) from night timepoints (when lights were off) (**Fig. 1d**) demonstrating that PC1 corresponds largely to day-night differences in activity preferences (**Fig. 1f**). PC2 represented variation in time points associated with changing light conditions (dawn and dusk, **Fig. 1d**), revealing the crepuscular (dawn/dusk) preferences of activity (**Fig. 1f**). PCs 3-10 had smaller contributions over the 24 hour period explaining only 0.5-7.5% of the variation in swimming speed. This analysis revealed that a species’ daily activity patterns have a diurnal/nocturnal component (PC1) and a crepuscular (PC2) component, and a species’ temporal activity pattern can be measured by their loadings for each. For example, hierarchical clustering based on the species loadings of PC1 and PC2 showed that the species form three main groups, which we designated as diurnal (negative PC1 loadings), crepuscular (high PC2 loadings, PC1 loadings near zero) and nocturnal (positive PC1 loadings) (**Fig. 1g**). In addition, several species exhibited high variability in activity patterns in our experiment (**Extended Data Fig. 1**). This included species such as *E. melanogenys* (Enamel) and *N. cygnatus* (Neocyg), some of which also displayed weak preferences for diurnal, nocturnal, and crepuscular periods (PC1 and PC2 loadings near 0). These species likely lack temporal preferences for activity and occupy a cathemeral lifestyle. Therefore, we added a cathemeral class to species which had very high variability (**Fig. 1g**, see methods). Interestingly, this analysis shows that a species’ preference for diurnal or nocturnal activity is not mutually exclusive with its preference for crepuscular peaks of activity. For example, *Tropheus moorii* (Tromoo) is diurnal (negative PC1 loading), but also crepuscular (positive PC2 loading), whereas *Astatotilapia burtoni* (Astbur) is diurnal (negative PC1 loading), but lacks crepuscular peaks of activity (low PC2 loading) (**Fig. 1g-h**). Together these results suggest that the Tanganyikan cichlids display all known activity patterns (diurnal, nocturnal, crepuscular, cathemeral). In addition, our results suggest that crepuscularity is not mutually exclusive with diurnality or nocturnality.

### Total rest is highly diverse in Tanganyikan cichlids

Given the diversity in the timing of activity across species, we next asked whether cichlids display variation in their total amounts of activity or inactivity per day. Inactivity can be used as a proxy for sleep in fishes; for example, it is commonly used in the diurnal zebrafish, which exhibit short bouts of reduced motility associated with higher arousal thresholds predominantly during the night^17,21,22^. Cichlids also displayed periods of inactivity. We used our high resolution tracking data to quantify the total daily amount of consistent inactivity periods (less than 5% of movement in a sliding 60s window), which we called “rest” (**Fig. 1i, see methods**). The majority of species rested near the substrate in the bottom quarter of the tank, including *E. melanogenys* (Enamel) and *Telmatochromis sp. “Lufubu”* (Telluf) (**Extended Data Fig. 2a-b**). Some exceptions were those with more pelagic lifestyles such as *Cyprichromis coloratus* (Cypcol), which tended to rest within the water column, and *Paracyprichromis nigripinnis* (Pcynig), which tend to rest off the bottom but next to the mesh walls of the arena (**Extended Data Fig. 2c**), with both species adopting a more vertical posture during rest compared to active periods (**Extended Data Fig. 2d**). We found extensive variation in total rest between species (**Fig. 1i**). For example, some species rested for up to 18 hours per day (e.g. *Altolamprologus sp.* “compressiceps shell” [Altshe]), while others displayed less than 3 hours of rest per day (e.g. *Ophthalmotilapia boops* [Ophboo] and *Limnotilapia dardennii* [Limdar]). These results demonstrated that the range in daily total rest in cichlids resembles the range known across all mammal species (∼90 million years of evolution), with bats resting for 20 hours per day and horses for only 3 hours^23^.

### Extensive temporal niche partitioning in Tanganyikan cichlids

We next asked whether differences in temporal activity patterns or total amounts of rest are associated with trophic ecology and diet, or morphology. To test this, we correlated our measured activity patterns (PC1 and PC2 loadings, and total rest) with a range of eco-morphological traits and diet guilds of these species^5^. We found that closely related species did not necessarily have similar activity patterns or amounts of total rest (**Fig. 2a**). No strong phylogenetic signal was observed for diurnal-nocturnal preference (PC1 loadings, Pagel’s 𝞴 = 0.630, Bloomberg’s K = 0.473), crepuscularity (PC2 loadings, 𝞴 ∼0, K = 0.438), or total rest (𝞴 = 0.609, K = 0.525), and behaviours were distributed relatively evenly across tribes (**Fig. 2a**). For example, strongly nocturnal and diurnal species were observed within the tribes Lamprologini, Ectodini, and Tropheini. Crepuscularity was also observed widely and seen in Lamprologini, Limnochromini, Ectodini, Cyprichromini and Trophenini (**Fig. 2a**). The species with the most rest included *Altolamprologus* spp. (Altshe, and Altcom) and *Telmatochromis* spp. (Telshe, and Telvit), as well as the mud hole dwelling *L. signatus* (Lamsig) (**Fig. 2a**). Species of the Tropheini had the least rest, and a large diversity in total rest was observed in both Lamprologini and Ectodini (**Fig. 2a**). These results show that in Lake Tanganyikan cichlids both closely related species and highly divergent species have highly divergent activity patterns and total rest amounts.

**Figure 2.**
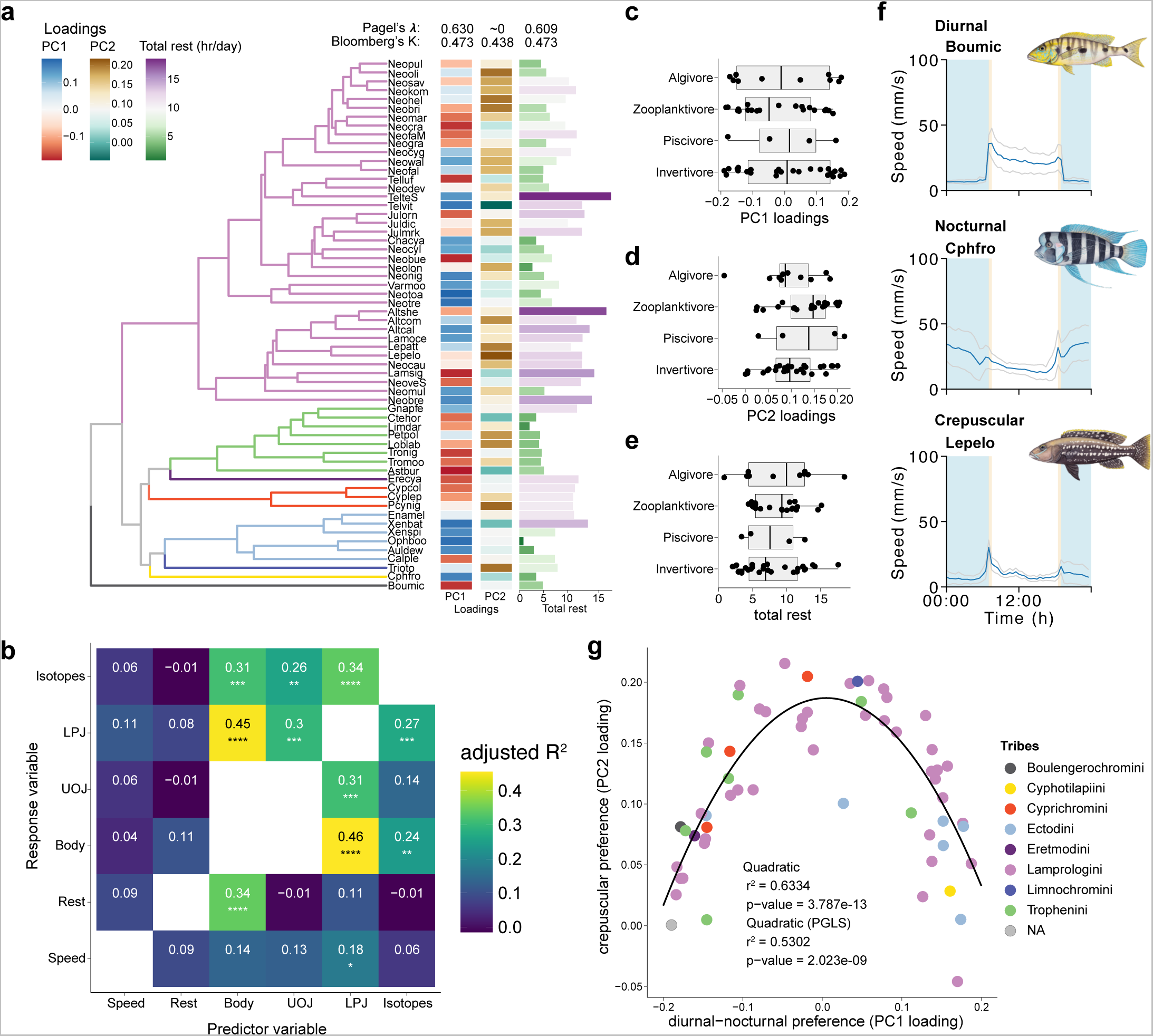
The weak relationships between behaviour and ecological measures supports temporal niche partitioning. **a,** The phylogenetic tree of the species in our study along with diurnal-nocturnal preference (PC1 loadings), crepuscular preference (PC2 loadings) and total rest, along with values for Pagel’s Lambda and Bloomberg’s K for each trait. **b,** Heatmap of adjusted R2 values for pairwise phylogenetically corrected two-block partial least squares analysis for comparisons between PC1 loadings, PC2 loadings, total rest and published data for stable isotopes values and datasets of body and jaw (UOJ: upper oral jaw; LPJ: lower pharyngeal jaw) morphology^5^. Numbers and colour represent R2 values; * = p-value <0.05, ** = p-value < 0.01, *** = p-value < 0.001. Diet guilds plotted against **c,** PC1 loadings, **d,** PC2 loadings, and **e,** total rest, each dot is one species. **f,** Examples of three species from the piscivore diet guild with diverse daily speed patterns, speed mean +/- SD. Species names are abbreviated using a six-letter code (**Supplementary Data 1**). **g,** The relationship between PC1 loadings and PC2 loadings can be best explained by a quadratic function, also when phylogenetically corrected. Species names are abbreviated using a six-letter code following Ronco *et al.* 2021 (**Supplementary Data 1**).

To test for a correlation between activity patterns and the environment occupied by a species, we compared previously generated stable isotope measurements and datasets of body and jaw morphology^5^ to our temporal activity patterns, using a phylogenetically corrected two-block partial least squares analysis (PLS). Stable isotopes measure a species relative position on the bentho-pelagic axis (δ^13^C) as well as their relative trophic level (δ^15^N). These data have previously been linked to morphological adaptations in cichlid body shape and oral and lower pharyngeal jaw morphology, which represent unique adaptations in feeding ecology^5^. As has previously been shown across the entire radiation(Ronco *et al.*, 2021), PLS scores for morphological traits strongly and significantly correlated with PLS scores of stable carbon and nitrogen isotope values across the species in our dataset as well (**Fig. 2b**). PLS scores for body shape were significantly correlated with total rest across the species in our dataset (**Fig. 2b**). In contrast, PLS scores representing diurnal, nocturnal, and crepuscular preferences of this study were not significantly correlated with either morphology or environment (**Fig. 2b**, **Extended Data Fig. 3**). These results reveal that temporal activity patterns in cichlids are not associated with specific morphologies or ecologies.

To investigate the associations between temporal activity preferences and total rest with diet guilds, we compared the behaviour of algivores, invertivores, piscivores, and zooplanktivores in our dataset. Diurnal-nocturnal preferences, crepuscularity, and differences in total rest were evenly distributed across diet guilds, and a phylogenetically corrected ANOVA revealed that there are no exclusive relationships between diet guild and activity pattern (**Fig. 2c-e, Extended Data Fig. 3**). For example, *B. microlepis* (Boumic), *C. frontosa* (Cypfro), and *L. elongatus* (Lepelo) are all medium to large piscivorous cichlids with wide distributions, but have diverse temporal activity patterns, including preferences for diurnal, nocturnal, and crepuscular periods, respectively (**Fig. 2f**). In contrast, the algivore *Tropheus sp.* “Black” (Tronig), the invertivore *N. buescheri* (Neobue), and the zooplanktivore *N. marunguensis* (Neomar) occupied diverse diet niches, but the same diurnal temporal niche (**Extended Data Fig. 4**). This analysis suggests that cichlids with similar habitat and diet niches can occupy all possible temporal niches, and that cichlids with similar temporal niches can occupy diverse habitat and diet niches.

### Temporal activity patterns in cichlids are part of a continuum bridged by crepuscular states

Our high resolution and extensive behavioural data allowed us to directly measure the relationship between diurnal-nocturnal preference (PC1 loading), crepuscularity (PC2 loading), and total rest. No significant correlations were observed between total rest and either diurnal-nocturnal preference or crepuscularity, with or without accounting for phylogenetic relatedness (**Extended Data Fig. 5**). Interestingly, though no linear correlation was observed between diurnal-nocturnal preference (PC1 loadings) and crepuscularity (PC2 loadings), they exhibited a clear parabolic relationship (**Fig. 2g**). Within our dataset, species with strong diurnal or nocturnal preferences (highly negative or positive PC1 loadings) had weaker crepuscular preferences (low PC2 loadings), and those with strong crepuscular preferences (high PC2 loadings) had weaker diurnal or nocturnal preferences (PC1 loadings near 0). This relationship was confirmed with a quadratic model, with PC1 loadings able to explain between 51-60% of variance in PC2 loadings (**Fig. 2g**). Cathemeral or crepuscular states have been suggested to provide a so-called ‘bridge’, facilitating adaptation of physiologies between dramatically different day and night environments^24–26^. The continuum of activity patterns in cichlids that we have observed suggests that an intermediate behavioural state with strong crepuscular preferences might have facilitated shifts between diurnal and nocturnal activity preferences in cichlids.

### The genetic signatures of activity patterns are complex and differ between clades

To investigate the genetic basis of temporal activity patterns and total rest, we followed an approach used in a recent study on the genetic basis of exploratory behaviour in Lake Tanganyika cichlids^27^. Briefly, we used a combination of a standard GWAS generalised linear model (GLM) and phylogenetic generalised least squares (pGLS) to account for phylogenetic relationships, and used cutoffs for genome-wide significance based on mutational simulations that should enrich for variants whose association with our behavioural traits is likely due to natural selection. Because of the relatively long history of the Tanganyikan cichlid radiation, and the deep ancestry of the species used in our study, recombination has broken down linkage between most variants in our dataset, and each single nucleotide polymorphisms (SNPs) is likely to provide an independent signature^27^. We used 120 whole genome sequences from 60 species to look for association between ∼39 million SNPs and (1) day-night preference (PC1 loadings), (2) preference for crepuscular activity (PC2 loadings), and (3) total amount of rest.

The above approach allowed us to identify 766 highly associated variants (HAVs) for day-night preference, 774 HAVs for crepuscularity, and 752 HAVs for total rest that passed our genome wide significance threshold (**Supplemental Data 3**, **Fig. 3 and Extended Data Fig. 6**). HAVs were distributed evenly across the genome, and no overlap was observed between HAVs for different behaviours (**Extended Data Fig. 7**). Individual HAVs demonstrated clear association with our behavioural traits, including HAV NC_031975:1851366, which was found in species with strong nocturnal preference across tribes (e.g. *N. tretocephalus* [Neotre], *N. toae* [Neotoa], *O. boops* [Ophboo], *Xenotilapia spilopterus* [Xenspi], and *X. bathyphilus* [Xenbat]), but not in those with strong diurnal preferences (**Fig. 3a**).

**Figure 3.**
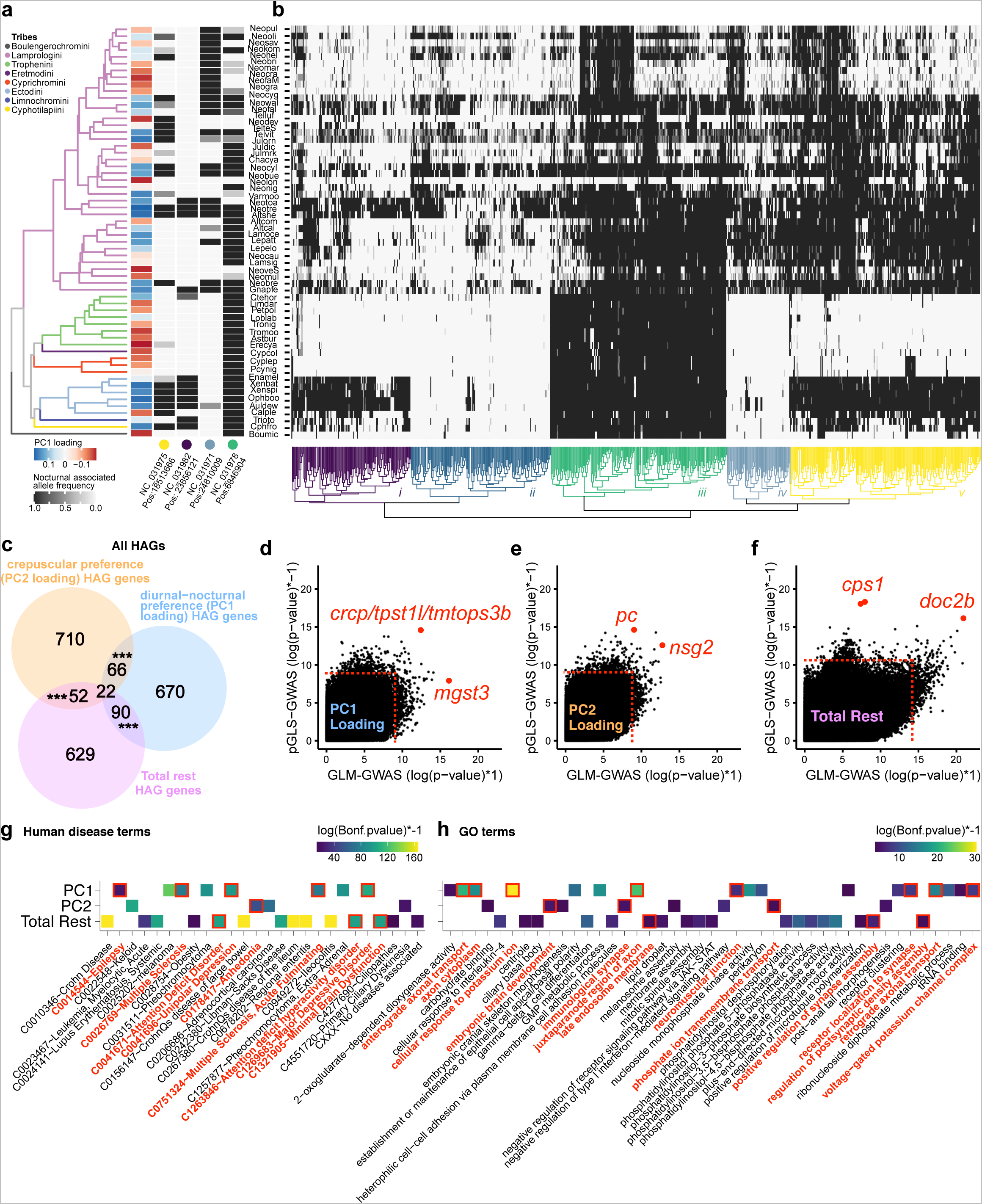
Genome wide signatures of cichlid temporal activity patterns. **a**, Phylogenetic tree of the species in our dataset along with the PC1 loadings (diurnal-nocturnal preference) and the allele frequencies of the high PC1 loading associated allele at four representative HAV loci. The coloured dot represents the cluster that each HAV is representative of from (b). **b**, Heatmap of the allele frequencies of the high PC1 loading associated allele across all highly associated variants (HAVs) for PC1 loading (diurnal-nocturnal preference). **c**, Venn diagram of the overlap of highly associated genes (HAGs) for PC1 loadings, PC2 loadings, and for total rest, annotated with the number of genes in each category. *** = p-value < 0.001 for enrichment tests between pairwise gene sets. **d-f**, Scatter plots of the pGLS-GWAS p-value and GLM-GWAS p-values for all SNPs associated with PC1 loadings (**d**), PC2 loadings (**e**), and total rest (**f**). The SNPs with the lowest p-value in the pGLS and GLM tests are labelled. Dotted lines indicated genome wide cutoffs for identification of HAVs. Tile-plots of human diseases (**g**), and GO terms (**h**) identified by gene ontology analysis with snp2go showing enrichment for the human orthologs of genes nearby to HAVs for PC1 loadings, PC2 loadings, and total rest. Tiles are coloured by the Bonferroni corrected p-value for the association test. GO categories associated with neuronal and synaptic function are highlighted in red.

To more systematically interrogate these patterns, we determined the directionality of the association between each allele and each behaviour, and clustered HAVs based on the frequency of whichever allele (reference or alternative) associated with positive PC1 loading scores (categories i-v, **Fig. 3b**), positive PC2 loading scores (**Extended Data Fig. 6a**), and high amounts of total rest (**Extended Data Fig. 6b**). For example, many nocturnal-associated alleles were mostly found within the Lamprologini (e.g. HAV NC_031971:2481009) (**Fig. 3a**), or mostly within the Ectodini (e.g. HAV NC_031982:23856121) (**Fig. 3a**). Additionally, there were alleles which were present in the majority of species, and whose absence was associated with strong diurnality, including *N. buescheri*, which is one of the most strongly diurnal species in our study (e.g. HAV NC_031978:6846904, **Fig. 3a**). Similar trends were observed for HAVs associated with crepuscularity and total rest, with alleles shared across the radiation, alleles specific to certain tribes (e.g. Lamprologini or Ectodini), as well as alleles whose absence was associated with the trait (**Extended Data Fig. 6**). Together, these analyses suggest that the genetics of temporal activity patterns in cichlids are complex, and that much of this complexity arises from differential association of HAVs between clades.

### Diel activity patterns are associated with synapse function and genes associated with neurological disorders, not the circadian clock

To identify putative molecular functions for HAVs associated with temporal activity patterns, we annotated all HAVs using SnpEff^28^. We termed such genes highly associated genes (HAGs)^28,29^. This approach identified 848 HAGs with diurnal-nocturnal preference, 850 HAGs for crepuscularity, and 793 HAGs for total rest. Unlike with HAVs themselves, we did observe an enrichment in the pairwise overlaps between HAGs for all three behaviours (**Fig. 3c**). The top HAVs for temporal activity patterns and total rest were associated with genes with known functions in the nervous system, including the regulation of sleep, or genes associated with neuronal disease and dysfunction in humans and model organisms.

For example, the most significantly associated variant to diurnal-nocturnal preference identified by pGLS-GWAS was within the intron of the gene *tpst1l* and downstream of both *tmtops3b* and *crcp* (**Fig. 3d**, **Extended data figure 7a**). *Teleost multiple tissue opsin 3b* (*tmtops3b*) is a non-visual opsin that may allow detection of blue light in hypothalamic deep brain nuclei^30^. Though not the closest gene to the HAV, *crcp* is the receptor for a neuropeptide (CGRP) with functions similar to hypocretin/orexin and PDF in regulating rest/wake in zebrafish and flies, respectively, and acts downstream of circadian pacemaking neurons^31,32^ (**Fig. 3d**). A SNP in the intron of *cacnb3b* was also strongly associated with diurnal-nocturnal preference. *cacnb3b* is a subunit of the voltage gated calcium channel, and another subunit of this channel (CACNA1C) has been linked to sleep latency and quality across multiple human populations^33,34^.

Two of the top HAVs for crepuscularity were a SNP in an intron of the pyruvate carboxylase gene (*pc*), and one downstream of the *nsg2* gene (**Fig. 3e**, **Extended data figure 7b**). Pyruvate carboxylase is involved in the production of neurotransmitters, and its deficiency is associated with both neurodevelopmental defects and seizures/epilepsy^35^. *Nsg2* is required for normal synapse maturation and regulates excitatory neurotransmission^36^. Mice lacking *nsg2* also display reduced activity at night suggesting it is involved in temporal activity preferences as well^37^. For total rest, our analysis also identified two adjacent SNPs separated by one base pair in the 5th to last intron of the carbamoyl-phosphate synthase 1 gene (*cps1*), and one SNP in the fourth intron of the gene encoding double C2-like domains beta (*doc2b*) (**Fig. 3f**, **Extended data figure 7c**). *Cps1* has been linked to diseases associated with lethargy^38^, and *doc2b* regulates spontaneous neurotransmitter release in the hippocampus^39,40^.

Enrichment analysis of HAGs for temporal activity patterns and total rest further implicated pathways and GO terms associated with synaptic transmission or assembly, and human neurological disorders, including attention deficit disorder (ADHD), epilepsy, and depression (**Fig. 3g-h**). Despite the importance of light and the internal circadian clock for cichlid activity patterns (**Extended Data** Fig. 8a-d)^14^, functions related to the regulation of the circadian clock, or melatonin regulation/signalling were not observed to be enriched in the HAGs (**Fig. 3c-f** and **Extended Data Figs. 7, 8e-g**). Together, these results suggest that evolutionary transitions between activity patterns, including diurnal to nocturnal transitions, are associated with genes regulating synaptic neurotransmission, rather than those involved in the circadian clock.

## DISCUSSION

Here, using the largest study of its kind, we characterised the temporal activity patterns of 60 ecologically diverse species of cichlid fishes from Lake Tanganyika. We provide evidence that temporal niche partitioning was an important component of niche diversification in this cichlid adaptive radiation. Our work extends on a study examining the activity patterns of 11 cichlid species over a 24 hr period from the Lake Malawi cichlid adaptive radiation^14^. In that study, while most species were diurnal or had no clear rhythm, one displayed nocturnal activity, prompting the authors to speculate that different activity patterns may have contributed to niche partitioning within the lake^14^. Our study provides extensive evidence of temporal niche partitioning by examining in depth (6 days/nights) a large number of species (60) from a well characterised adaptive radiation in a ‘common garden’ setup. Moreover, by integrating behavioural data with eco-morphological data for each species, we observe that cichlid species which occupy similar habitat and diet niches feature different temporal activity niches, and species with similar temporal niches differ in their habitat, morphology, and diet specialisations. These results demonstrate that temporal niche partitioning is an independent axis of diversification.

Temporal niche partitioning has mainly been observed in distantly related species^7,9,41^. For example, temporal niche partitioning was observed in large coastal sharks^8^, and in mammals^9,42^. Conversely, several studies have looked at the temporal activity patterns of closely-related, but non-sympatric *Drosophila* species, and have found that these species all have similar temporal patterns, but differ most in their amount of activity^43,44^. Detailed studies of more species and other adaptive radiations in ecologically-relevant contexts (within social groups such as in Lloyd *et al.,* 2023^45^, and more complex environments) as well as studies in the wild will be required to understand the full complexity of daily activity patterns.

Furthermore, our work suggests that transitions between diurnal and nocturnal activity patterns are facilitated through an intermediate crepuscular state or “bridge”. This observation confirms and extends previous categorical studies on the temporal activity patterns of skinks^46^, geckos^47^, and mammals^24–26,48^, in which direct evolutionary transitions between diurnal and nocturnal activity patterns are slower than transitions from crepuscular/cathemeral to diurnal and nocturnal patterns. Our study provides high resolution quantitative, rather than categorical, behavioural evidence for species occupying a continuum of behavioural states consistent with this bridge hypothesis. Ancestral reconstructions of temporal activity across a more complete cichlid phylogeny could provide evidence of the “bridge” state at ancestral nodes where transitions are predicted to have occurred, and determine the tempo and dynamics of temporal niche partitioning in this system.

Our study also provides an important set of candidate loci that may underlie specific temporal activity patterns. Temporal activity patterns in cichlids are polygenic, primarily due to differential usage of SNPs across clades. It has been suggested that polygenic trait architectures are better than simple architectures at promoting rapid and stable speciation in sympatry^49,50^. Interestingly, we found no evidence for the involvement of circadian clock genes in diurnal, nocturnal, and crepuscular preferences, or in total rest. This lack of associations with clock genes suggests that evolutionary transitions in activity patterns occur independently of, and downstream of the core circadian clock. This is consistent with work showing that diurnal and nocturnal mammals have no systematic differences in their circadian timing systems^51,52^. Modelling studies have also demonstrated that changes in the activity phase of circuits downstream of the clock can explain switches between diurnal and nocturnal behaviour^53^. Indeed, we observed enrichment for genes associated with neuronal function and synaptic transmission. Many of these genes are also associated with human neurological and neuropsychiatric disorders, including Alzheimer’s, ADHD, and epilepsy/seizures. Furthermore, sleep phenotypes and neurological disorders have overlapping genetic components in humans^54^, and neuropeptides controlling sleep and wake states have been linked to the occurrence and severity of seizures^55^. Our results also suggest an overlap between the molecular mechanisms underlying evolutionary transitions in activity patterns and sleep duration and quality in humans from GWAS studies^33,34^. Future work will determine which SNPs and genes are causally linked to evolutionary shifts in activity patterns in cichlids, and whether these are the same mechanisms governing shifts across other clades, including mammals.

## METHODS

### Fish husbandry

All animal work was performed at the facilities of the University of Basel. The initial screen of 60 cichlids species was performed at the Zoological Institute (University of Basel). Generally, all species in our study were housed in similar conditions (12 hours of light, 12 hours of darkness; 24-25℃ water temperature) with 30 minute ramping light conditions at dawn and dusk. Cichlids were fed regularly, with fortnightly water changes. Dark/dark experiments were performed at the Biozentrum (University of Basel). Here, species were housed in a recirculating system (Tecniplast) with 8% exchange of water everyday with 14 hours of light, 10 hours of darkness; 26℃ water temperature with 15 minute ramping light conditions at dawn and dusk. Fish were fed twice a day, once with live food and once with dry food.

Cichlids were kept at densities no greater than 0.5 cm/litre for fish up to 5 cm total length and 1 cm/litre of water for larger fish up to 15 cm total length, and generally kept in species-specific tanks. Environmental enrichment varied between species according to their ecology, but generally consisted of rocks and terracotta plant pots serving as refuge and breeding areas, sandy or gravel substrates, and shells. The source of animals varied, but they stem generally from in-house natural breeding of populations originally collected in Lake Tanganyika, or purchased from local suppliers (Garten-und Zoobedarf Schrepfer). All experiments were performed under holding permit nrs. 1010H and 1035H, and experimental permit nrs. 2356 and 3102 issued by the cantonal veterinary office Basel.

### Behavioural assays

We developed a behavioural assay based on a previous study^19^, to record daily activity patterns of individual fish. Glass tanks (45cm height x 110cm length x 25cm depth, clear glass on one long side and opaque glass on 3 sides, Pavlica Akvária) were used to house fish for experiments. Each tank had a thin layer of sand and was physically divided into arenas for individual fish. For most species the tanks were divided into four arenas (each 25cm wide), however, for the larger fish tanks were divided into larger arenas. For *Boulengerochromis microlepis* (Boumic) juveniles tanks were split into three arenas (33cm wide). For *Enantiopus melanogenys* (Enamel) and the larger individuals of *Limnotilapia dardennii* (Limdar), tanks were split into two arenas (50cm wide). The dividers were made of PMMA opal white with a mesh insert allowing for water flow and visual communication of fish to their neighbours. Fish were kept on a 12h:12h light:dark cycle from 07:00 to 19:00 CET with the light ramping on from 07:00-07.30 and ramping down from 18.30-19:00 (TC420 light controller). With the exception of the species used in the dark:dark experiments, which were housed on a 14h:10h light:dark cycle from 08:00 to 22:00 with the light ramping on from 08:00-08.30 and ramping down from 21:30-22:00. This was to account for the fish being kept in a different aquarium room with a 14:10h cycle instead of the 12:12h cycle. The tanks were backlit with a panel of infrared LED lights which were diffused by the opaque glass and a diffuser to generate even lighting and allow recording during the night.

Using this setup we assayed 60 species of cichlid fishes from Lake Tanganyika (**Fig 1a**). To increase the number of species we could test we used cichlids from a range of sources including wild caught and aquarium bred fish. We aimed to assay 7-12 fish per species, but in some cases we included rarer species for which we only had less than 7 individuals (*Chalinochromis cyanophleps* [Chacya], *Lobochilotes labiatus* [Loblab]: 2 individuals; *Altolamprologus compressiceps* [Altcom]: 3 individuals; *Neolamprologus walteri* [Neowal]: 4 individuals; *Eretmodus cyanostictus* [Eracya], *N. falcicula* [Neofal], *N. sp. ‘Kombe’* [Neokom], *Xenotilapia spilopterus* [Xenspi]: 6 individuals; see **Supplementary Data 1**). Fish were fed according to their home tank schedule of every couple of days as determined by the fish technicians. Because the fish and the behavioural setup were kept in a room where the temperature could not be well controlled, temperature varied depending on season, and was generally between 24 to 28℃. Animals which became sick, died or escaped were excluded. Age of tested fish varied, almost all fish were tested when they had indicators of sexual maturity (colouration, behaviours, breeding), however, *B. microlepis* (Boumic) and *Cyphotilapia frontosa* (Cypfro) were tested as juveniles due to the long life cycles and lack of availability of adults of these larger species. Both male and females were tested, and sex of was recorded and used to disaggregate results when it could be reliably be determined.

### Processing of behavioural data

Each arena was recorded from the front using cameras (RoHS 1.3MP B&W Chameleon USB 3.0 Camera 1/3” CCD CS-Mount -CM3-U3-13S2M-CS, Flir) fitted with lenses (YV4.3x2.8SA-2, Fujinon) and long-pass filters to exclude white light (Midopt Near-IR Longpass slipmount Filter 780-30.5). Recording was done by custom Python3 v3.7 code utilising the Spinnaker SDK API, with processing from the following packages: numpy^56^, imageio^57^, opencv-python^58^, PyYAML, pandas^59^ and matplotlib^60^. Each arena was selected as a region of interest (ROI), and a 10fps mp4 video of this ROI was saved every hour. We tracked animals online using background subtraction and thresholding with a minimum object area of 100 pixels. Backgrounds were calculated off the 95th percentile of the previous 1hr movie. Positions of fish were interpolated during frames where the fish was not visible (e.g. if it was hidden in a corner, or behind sand). Timestamps, X and Y position and surface area of the tracked object were saved alongside each video as a comma-separated file (csv), as well as a YAML file of metadata, using PyYAML v5.3.1 software^61^. Fish were recorded for a week, but only data from midnight on the first day to midnight on the second-last day was used for further analysis (6 days/nights total).

Extensive quality control was performed on each video to ensure accuracy of behavioural tracks across cichlid species. This primarily involved excluding portions where the tanks were briefly obstructed by personnel, or increasing background subtraction to 30, 20, or 10 minute periods (instead of 1 hour) to account for sand displacement by individual fish. Additionally, many videos were re-tracked with smaller ROIs to remove problematic regions associated with the edges of the original ROIs chosen. This included removal of pixels covering adjacent arenas (which can permit detection of adjacent fish), or pixels covering the top of the tank, which could contain disturbances in the water due to feeding or water exchange (water bubbles).

Swimming speed was calculated using subsequent X and Y positions. Video frames where speed exceeded a threshold of 200 pixels were replaced with the average of −/+5 frames (1 sec). Such high speeds were never observed for individual fish, but represented jumps caused by the background subtraction detecting objects elsewhere in the tank (water bubbles, or movement in adjacent arenas). Speed as well as X and Y position were smoothed by 0.5 sec windows. For most plots data is binned by 30 minutes (**Supplementary Data 2**).

### Exploration of behavioural data

To investigate the patterns of daily activity we used Principal Component Analysis (PCA). Averages of the daily speed per species in 30 minute bins were standardised by z-scoring (48 dimensions). PCA was run with 10 components using the PCA function from the sklearn.decomposition v0.0 package^62^. Ward clustering of PC1 and PC2 loadings separated the species into three groups, which based on the patterns we defined as diurnal, nocturnal and crepuscular. However, as we ran PCA on the daily averages it does not take individual variability into account, and we saw species with variable patterns, we therefore designated any species where the min to max daily average speed difference was smaller than two mean standard deviations, to be cathemeral (see **Fig. 1g**).

Besides speed, we also transformed activity into a measure of *rest*. First, movement was determined by a threshold of 15mm/s, above this threshold the animal is actively moving, while below this threshold most activity is noise (artefacts of tracking, or small undulations due to fin movements). To uncover behavioural states where the fish was mostly inactive, we used *rest*. A time point was set to be positive for rest when there was less than 5% movement in a sliding 60s window. Like speed, rest was also binned in 30min bins. We also quantified the position of fish, here we scaled the data between 0 and 1, where 0 was the minimum and 1 was the maximum fish position. Paired t-tests with Bonferroni correction were used to test for significant differences for vertical rest position between active (non-rest) and rest states.

To find rhythmicity in the speed data we used periodograms. We calculated these using code from the CosinorPy package v3.0 using the 30min binned speed data^63^. The default CosinorPy threshold (0.05) was used to identify significant spectra peaks.

Plots were made using the following software packages: Matplotlib^60^, Seaborn^64^, CMasher^65^.

### Comparing behaviour to ecological features

Eco-morphological data for all cichlids species in our study were taken from Ronco et. al.^5^. Three of our focal species are not included in the Ronco *et. al.* dataset as they are either notpart of the radiation, or are not found within the lake. These excluded species are: *A. burtoni* (Astbur), *N. devosi* (Neodev), and *Telmatochromis sp. “lufubu”* (Telluf). We used data on body, upper oral jaw, and lower pharyngeal jaw morphology, and stable isotope values (δ^13^C, δ^15^N). We compared these values to the metrics derived above, specifically diurnal-nocturnal preference (PC1 loadings), crepuscular preference (PC2 loadings), and total rest per species.

Phylogenetic signal in temporal activity pattern traits (PC1 and PC2 loadings, total rest) was tested using the function phylosig from the package phytools^66^. To investigate the links between eco-morphological traits and temporal activity patterns we performed pairwise phylogenetically corrected two-block partial least squares analyses, alternating each trait between predictor and response variables using the R packages pls^67^ and geomorph^68,69^. Importantly, PC1 and PC2 for temporal activity patterns which were calculated by PLS represented day-night preference and crepuscularity, and were highly similar to principal components for activity patterns (PC1 and PC2 in **Fig. 1** above). To investigate links between cichlid diet guilds and temporal activity patterns, cichlids were grouped by diet and a phylogenetic ANOVA was performed using the function aov.phylo from the geiger package^70^.

### Identifying genetic variants associated with temporal activity patterns

Whole genome sequences were obtained from GenBank, and the accession numbers for the samples used are available in **Supplemental Data 4**^5^. This dataset contains two individuals (one female and one male) from each species included in our behavioural analysis. Therefore, whole genome sequences from 60 species were used to generate a new variant call set specific to this study based on alignment to the *Oreochromis niloticus* genome (O_niloticus_UMD_NMBU, GCF_001858045.2, NCBI). We followed GATK best practices and analysis pipelines to align and call variants (GATK version 4.2.4.0), and used genome masks generated with custom scripts for variant filtration. This was followed by association studies to identify highly associated variants (HAVs), and analysis of genes nearby to HAVs with custom R scripts. All steps are outlined in more detail in the following sections.

### Alignment, variant calling, and variant filtration

Short reads from each individual fish were processed using MarkIlluminaAdaptors before being aligned to the most recent *O. niloticus* genome assembly (O_niloticus_UMD_NMBU, GCF_001858045.2, NCBI) using bwa mem software (version 0.7.17)^71^). Aligned reads were processed using MergeBamAlignment, SortSam, and MarkDuplicates in picard (version 2.26.2)^72^. Single nucleotide polymorphisms (SNPs) and short insertions and deletions were detected against the reference genome for each individual using HaplotypeCaller across 80 genomic intervals of equal length. Variant calls across genomic intervals were then combined using GatherVcfs, and the resulting variant call format (vcf) files for each individual were collected using CombineGVCFs. All samples were then jointly genotyped using GenotypeGVCFs resulting in a single vcf file containing variant calls across all sites and for all 60 species.

We performed extensive filtering on the genotyped variant file using three separate genome masks. The first genome mask was generated by identifying low quality sites using the VariantFiltration function and the expression “QD < 2.0 || FS > 60.0 || MQ < 40.0 || MQRankSum < −12.5 || ReadPosRankSum < −8.0” in GATK. The second genome mask identified variants whose read depth was not less than 900 or greater than 1900 when summed across all samples. Cutoffs for read depth were determined by examining the distribution of read depth from a random subset (10%) of genotyped sites and identifying a region with roughly normal distribution. The third genome mask was based on the ability of pseudo-reads generated from the *O. niloticus* genome to reliably map back to the correct location. We used the SNPable tool to divide the reference genome into overlapping 100 k-mer sequences (http://lh3lh3.users.sourceforge.net/snpable.shtml), and to generate an intermediate mask. This mask was then converted into bed format using a modified version of the makeMappabilityMask python script from msmc tools (https://github.com/stschiff/msmc-tools/tree/master). All three masks were then merged and used to hard filter genotype variants using VariantFiltration and SelectVariants.

This list of variants that passed the above masking approach was subjected to one final filtering step to simplify association studies. We excluded sites where the minor allele was present in only a single individual, all multiallelic sites, sites with insertions or deletions, and those that mapped to unplaced scaffolds. This filtering pipeline resulted in the identification and selection of roughly 39 million SNPs.

### Genome-wide association of variants to temporal activity patterns

To associate variants with temporal activity pattern preferences and total rest we first estimated allele frequencies for each SNP using the evo software package v.0.1 r23 and the subprogram alleleFreq following the approach of Sommer-Trembo *et al.,* 2024^27^ (https://github.com/millanek/evo). Briefly, we used genotype likelihoods from GATK, and assumed a Hardy-Weinberg prior to obtain posterior probabilities for reference and alternative allele frequencies at each loci. Allele frequencies derived using this approach were then used to test for associations between temporal activity pattern preferences and total rest.

We used a combination of a general linear modelling (GLM) and phylogenetic generalised least squares (pGLS) to identify associated variants, which accounts for phylogenetic relationships, as well as the possibility of allele-sharing between species. In association tests, diurnal-nocturnal preference (PC1 loadings), crepuscular preference (PC2) and total rest were used as response variables, and linearly scaled allele frequencies (ranging between −1 and 1) at each of the 40 million SNPs as the predictor. Both models were run in the R environment (version 4.0.3). The GLM was run using the command lm(temporal activity phenotype ∼ allele frequency), and iterated over each SNP. For pGLS we used the caper package^73^ and the function pGLS using the command pGLS(temporal activity phenotype ∼ allele frequency, phylogenetic tree), where the phylogenetic tree was that of the Lake Tanganyika cichlids from Ronco *et al.,* 2021^5^ pruned to include only the focal species in our study.

### Selection and analysis of highly associated variants (HAVs)

We next wanted to identify SNPs whose allele frequencies were strongly or significantly associated with temporal activity preferences and total rest. Our study is potentially underpowered to identify significant associations between traits and allele frequencies using multiple-testing correction, due to the low number of species (60), and the high number of tested alleles (∼40 million). However, a previous study has suggested that selecting the top fraction of SNPs that correspond to the lowest p-values in GLM and pGLS tests enriches for associations that are unlikely due to natural processes in the absence of selection^27^. Accordingly, we decided to focus on the top 99.99th percentile of SNPs with the lowest p-values for each association test (GLM and pGLS) and for each trait. This resulted in 766 unique SNPs that were highly associated with diurnal-nocturnal preference (PC1 loadings), 774 for crepuscular preference (PC2 loadings), and 752 for total rest. For each highly associated variant (HAV), we determined which allele (reference or alternative) was associated with nocturnal preferences (high PC1 loadings), crepuscular preference (high PC2 loadings), or total amounts of rest using the coefficients of the GLM models in R. These were then used for plotting allele frequencies, and to determine clusters of SNPs with distinct patterns across our species using hierarchical clustering and the function hclust in R.

### Gene ontological analysis of genes associated with HAVs

We next sought to identify genes that might be in proximity to the identified highly associated variants, and therefore may underlie temporal activity preferences or total rest. To annotate each SNP we used snpEff (version 5.2)^28^ and a genome database built from the *O. niloticus* UMD NMBU genome assembly downloaded from the NCBI RefSeq database. Human orthologs for *O. niloticus* genes were identified using the NCBI datasets command line tools (https://github.com/ncbi/datasets), as well using Ensembl biomart (Ensembl release 111). To identify gene categories that were enriched we used the software SNP2GO, which annotates all SNPs with GO terms based on their proximity to genes, and therefore allows testing the enrichment for SNPs that affect particular GO terms. We used modified versions of the function snp2go, which allowed the testing of disease and tissue expression associated gene sets collected using the packages PhenoExam (version 0.1)^74^ and RDAVIDWebService (version 1.28.0)^75^ in R.

## Data and code availability

Scripts for online recording and tracking were written in python and are available online (https://github.com/annnic/cichlid-tracking). All scripts for performing quality control and running analysis of behavioural activity, including generation of plots of cichlid weekly and daily speeds were written in python and are available online (https://github.com/annnic/cichlid-analysis). Scripts for analysis of eco-morphological data, construction of phylogenetic plots, highly associated variant analysis, and gene ontology analysis were written in R and available online (https://github.com/maxshafer/cichlid_sleep_gwas). Scripts for running genome wide association analysis, including the GATK python, generation of genome masks, and variant identification and filtering were written in bash and available online (https://github.com/maxshafer/cichlid_sleep_gwas). The time-calibrated species tree, morphology and stable carbon (C) and nitrogen and (N) isotope signatures were taken from Ronco *et al.* 2021^5^ (data available on Dryad: https://datadryad.org/stash/dataset/doi:10.5061/dryad.9w0vt4bbf).

## ACKNOWLEDGEMENTS

We thank members of the Shafer, Schier and Salzburger laboratories for discussion and advice and A. Kempf and V. Bitsikas for valuable comments on this manuscript. We thank the Biozentrum mechanical and electrical workshops as well as D. Lüscher for technical support, J. Johnson for fish illustrations and K. Ntemos at University of Basel, sciCORE for discussions. This work was supported by grants from the Swiss National Science Foundation (SNSF) to M.E.R.S. (196313), A.F.S. (197837), M.M. (193464) and W.S. (208002), from the Human Frontier Science Program (HFSP) to A.L.A.N (LT000400/2019-L), and by the German Science Foundation (DFG; fellowship SO 1737/1-1) and the Research Fund for Junior Researchers of the University of Basel to C.S.-T.. Computing resources and infrastructure were provided by sciCORE (https://scicore.unibas.ch/), the Center for Scientific Computing at the University of Basel, with support from the Swiss Institute of Bioinformatics (SIB).

## AUTHOR CONTRIBUTIONS

A.L.A.N., M.E.R.S., W.S., and A.F.S. conceived and designed the study. A.L.A.N. and M.E.R.S. collected and analysed behavioural data, performed ecological comparisons, and the genomic analysis. A.I., A.R., R.G.-D., and L.F aided in study design and collection and interpretation of behavioural data. M.M. and C.S.-T. provided scripts and aided genomic analysis. A.L.A.N., M.E.R.S., W.S., and A.F.S. interpreted the results and wrote the manuscript. All authors read and approved the manuscript.

---

**Extended data Figure 1.**
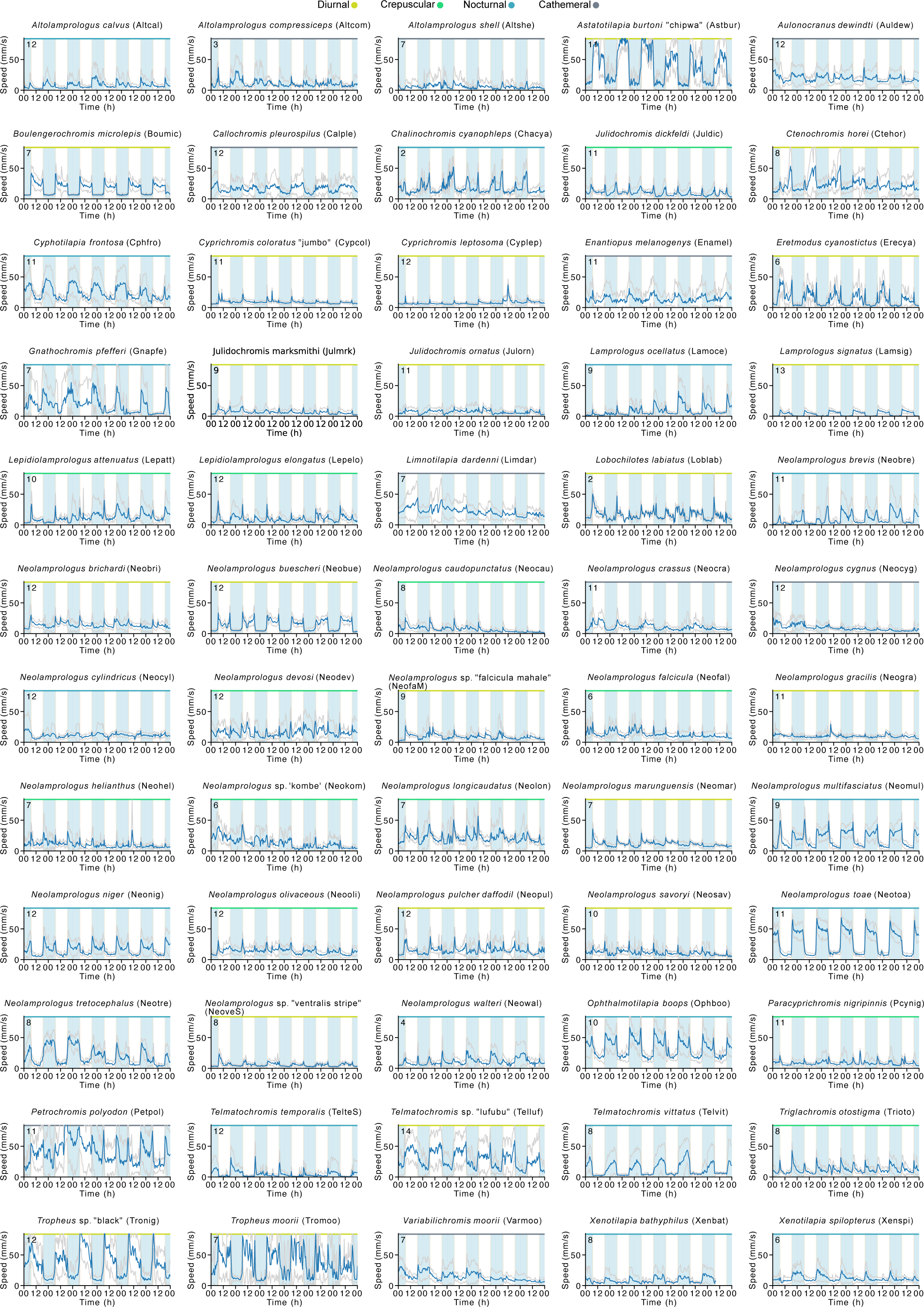
**a,** Weekly speed (mean +/- SD) traces for the 60 cichlid species assayed. Timeline and light cycle (12:12h light:dark) is the same as in the schematic shown in Figure 1b. Number of animals assayed shown in the top left corner. Temporal guild shown by top colour bar.

**Extended data Figure 2.**
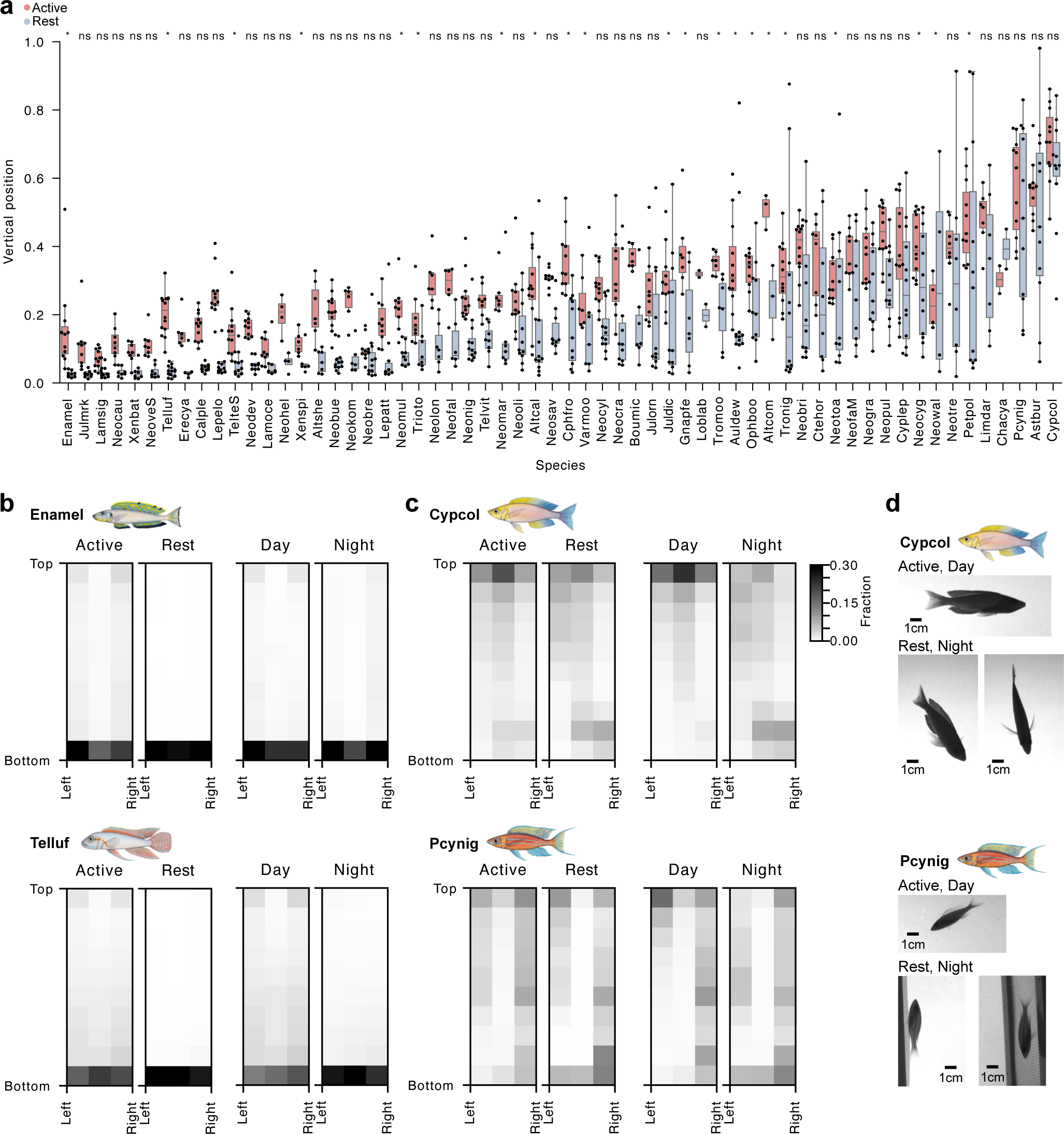
Many species prefer to rest at the bottom of the tank. **a,** Average vertical position (scaled: 0: bottom, 1: top) during rest or active periods for each species, ordered by lowest vertical position during rest. For the majority of the cichlid species, rest bouts occurred predominantly when at the bottom of the tank, either just above, or in contact with the sandy substrate. Significance level of the Bonferroni corrected p-values for paired t-tests of differences between mean vertical position of rest and active bouts are shown on top; * = p-value <0.05. Each dot shows the average for one individual. **b-c,** Density plots of positions for *Enantiopus melanogenys* (Enamel), *Telmatochromis sp. “Lufubu”* (Telluf), *Cyprichromis coloratus* (Cypcol) and *Paracyprichromis nigripinnis* (Pcynig) separated by Active and Rest or by Day and Night. **d,** Representative images of fish postures during Active phases of the Day, or Rest phases of the Night, for Cypcol and Pcynig. Species names are abbreviated using a six-letter code following Ronco *et al.* 2021 (**Supplementary Data 1**).

**Extended data Figure 3.**
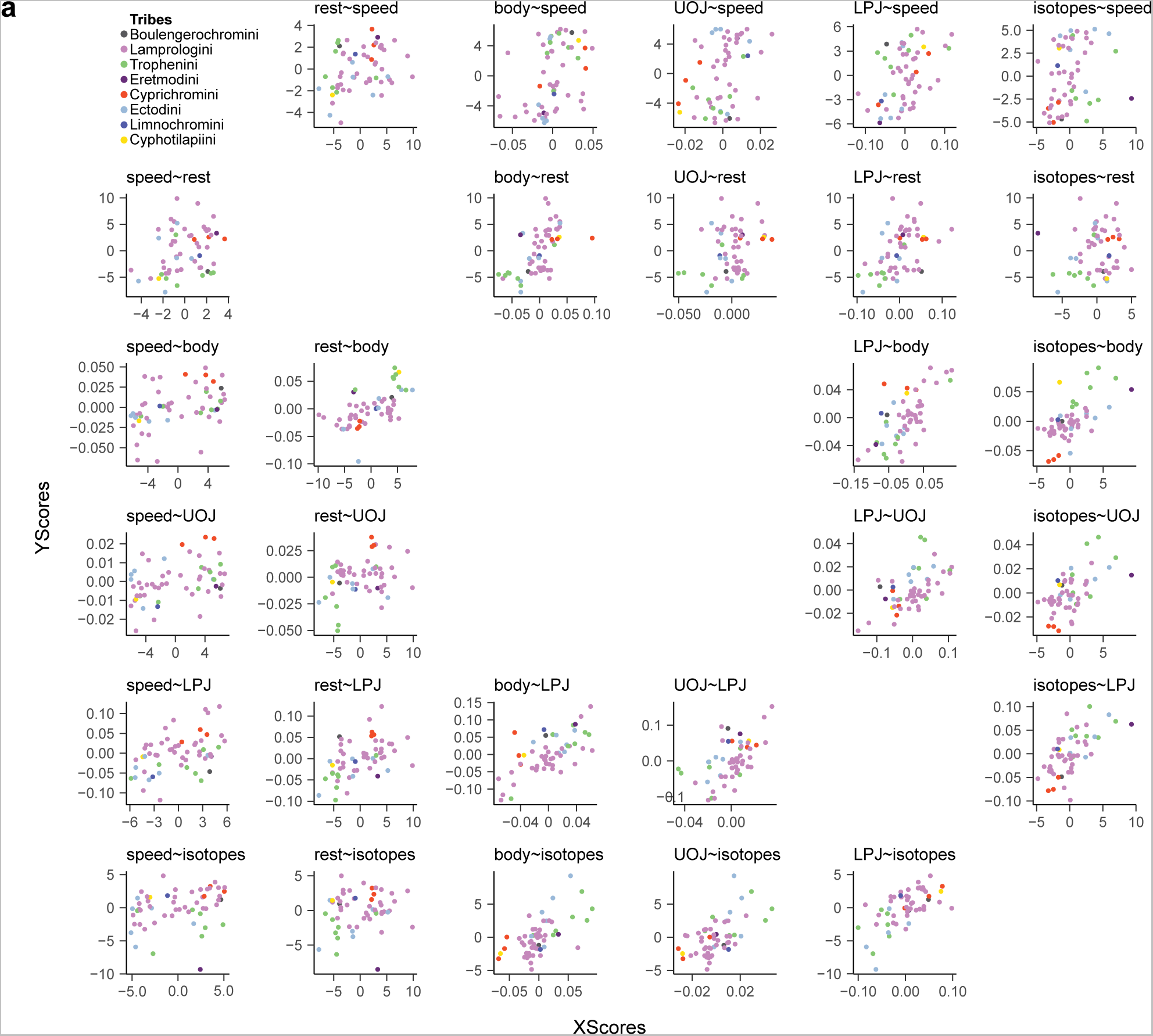
Relationships between temporal activity patterns and eco-morphological features of cichlids. **a**, Pairwise relationships between PC1 loadings, PC2 loadings, total rest and published data for stable isotopes values and datasets of body and jaw morphology (UOJ: upper oral jaw; LPJ: lower pharyngeal jaw)^5^. Dots represent each species and are coloured by their tribe.

**Extended data Figure 4.**
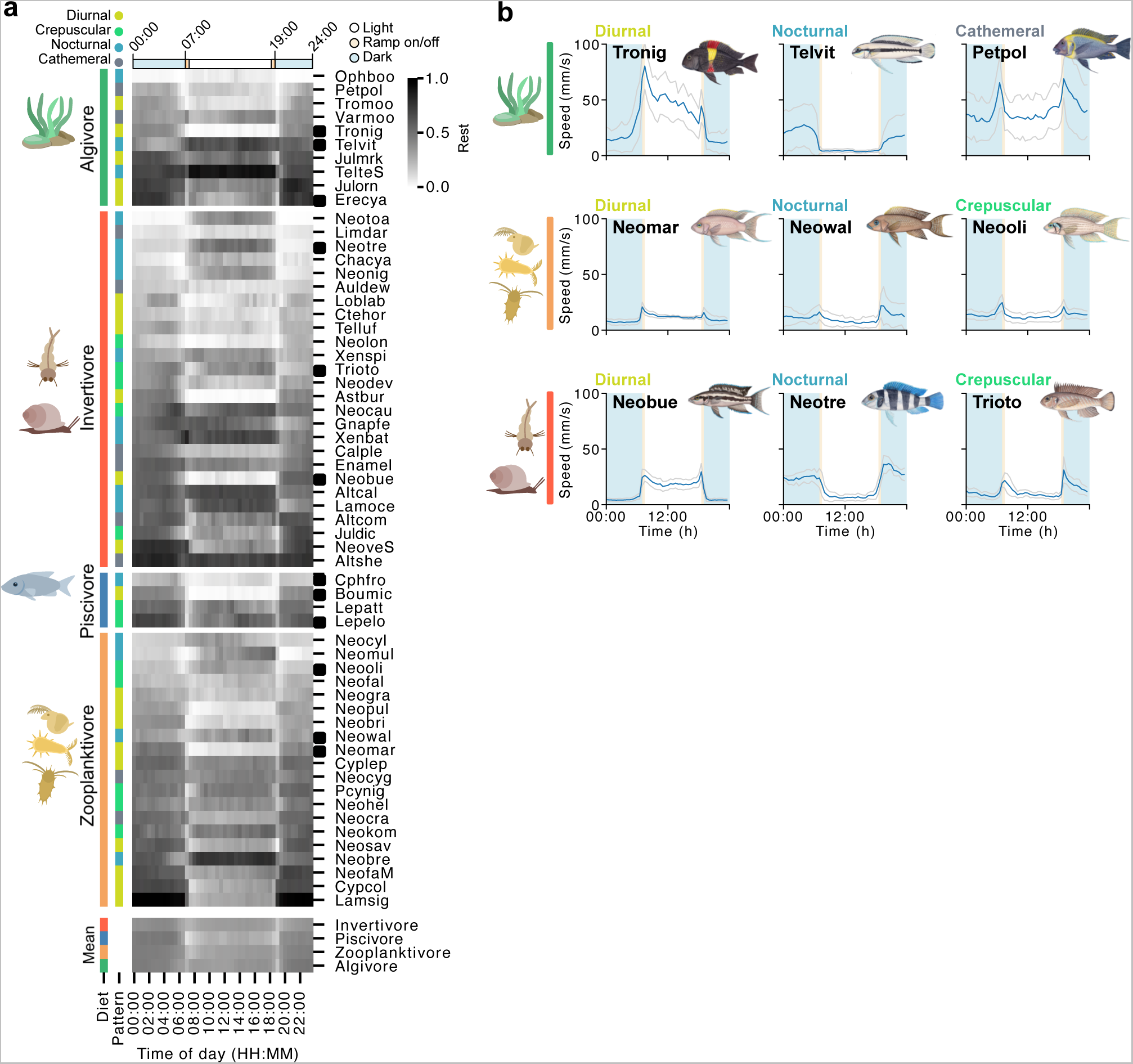
Ecological features and relationships to behavioural data. **a,** top: heatmap of daily average of rest for each species grouped by diet guild. bottom: mean rest for each group. **b**, Examples of three species with diverse daily speed patterns per diet guild: Algivore, Invertivore, and Zooplanktivore, speed mean +/- SD. Species names are abbreviated using a six-letter code following Ronco *et al.* 2021 (**Supplementary Data 1**).

**Extended data Figure 5.**
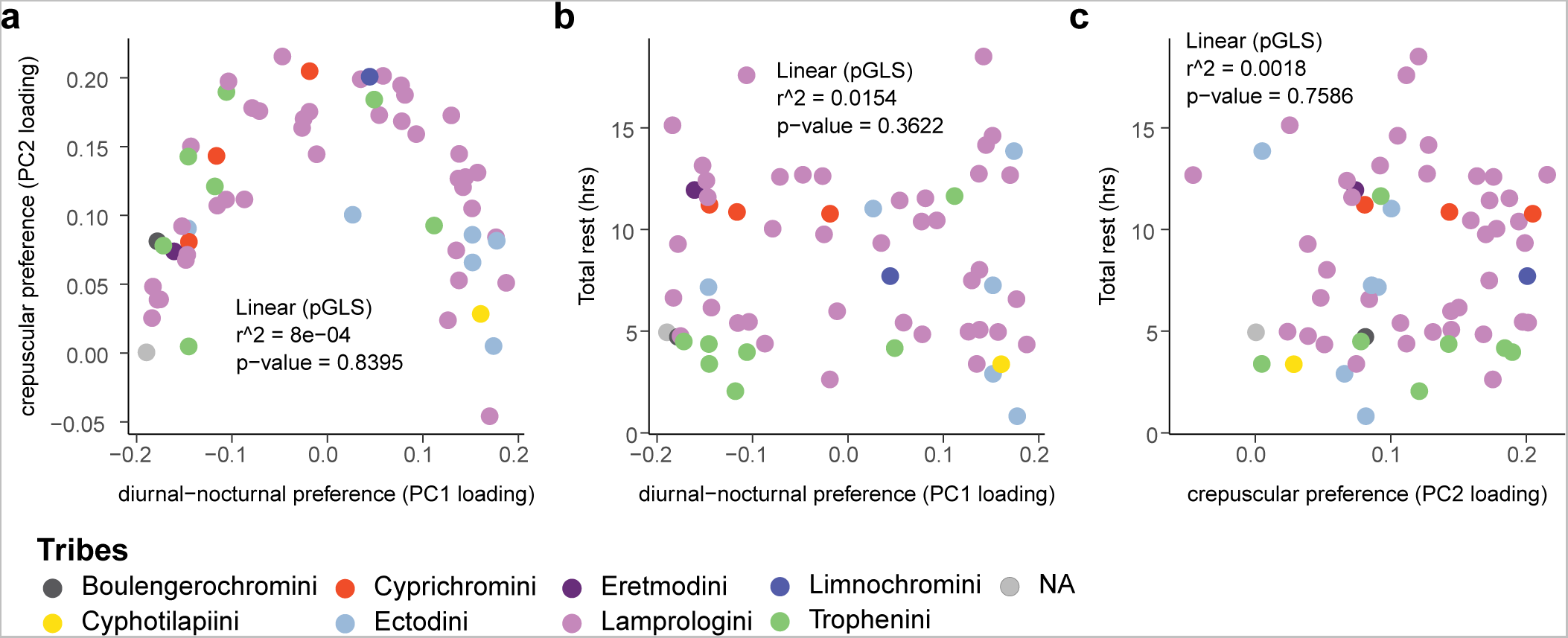
Relationships between temporal activity patterns. **a-c,** The phylogenetically corrected pairwise relationships between PC1 loadings, PC2 loadings, and total rest, along with the statistics for the fit of linear functions explaining the relationships. Dots represent each species and are coloured by their tribe.

**Extended data Figure 6.**
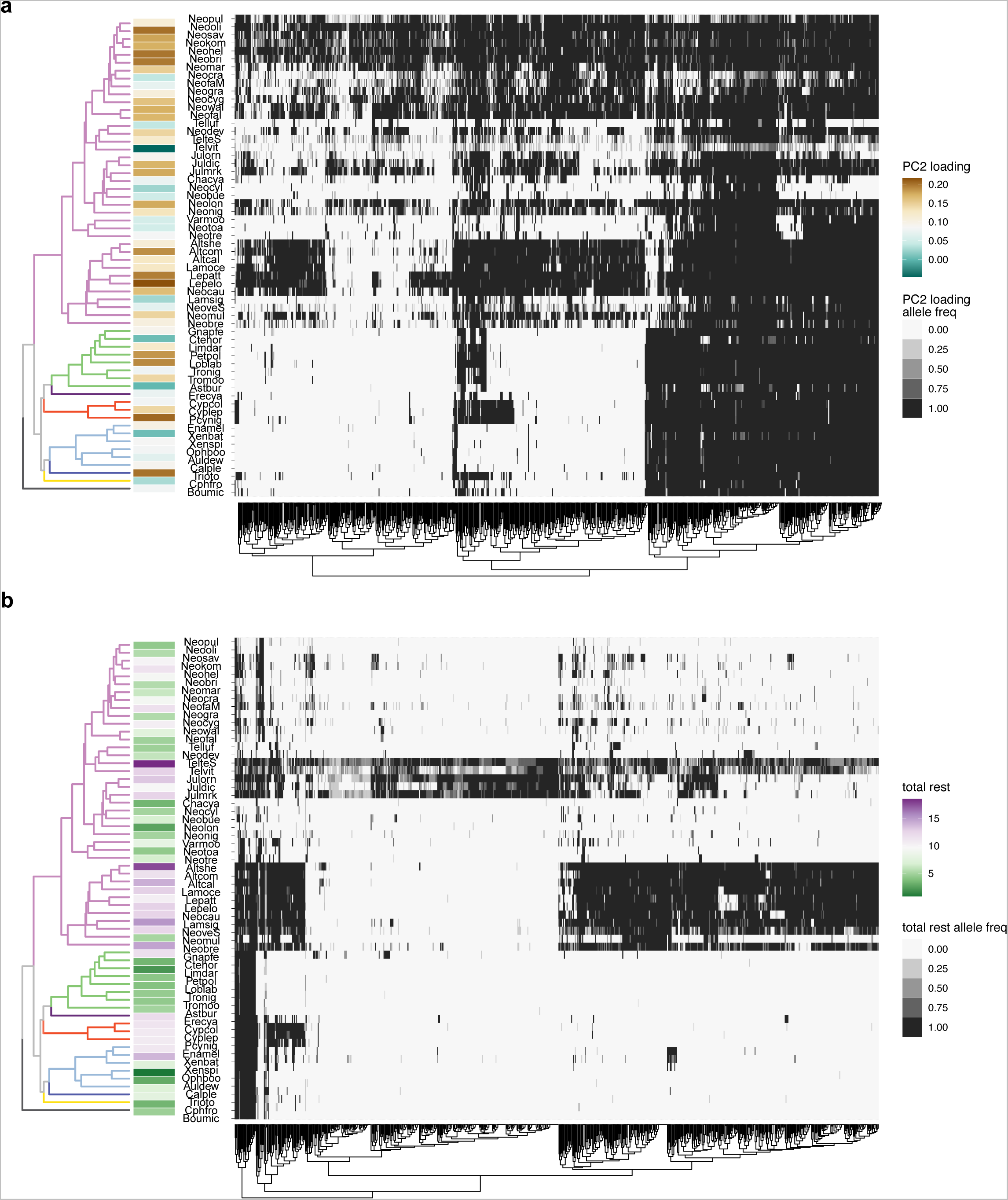
Clustering of highly associated variants for crepuscularity and total rest across cichlids. **a**, Phylogenetic tree of the species in our dataset along with a heatmap of the allele frequencies of the high PC2 loading associated allele across all highly associated variants (HAVs) for PC2 loading (crepuscular preference). **b**, Phylogenetic tree of the species in our dataset along with a heatmap of the allele frequencies of the high total rest associated allele across all highly associated variants (HAVs) for total rest. Dendrograms represent relationships between HAVs.

**Extended data Figure 7.**
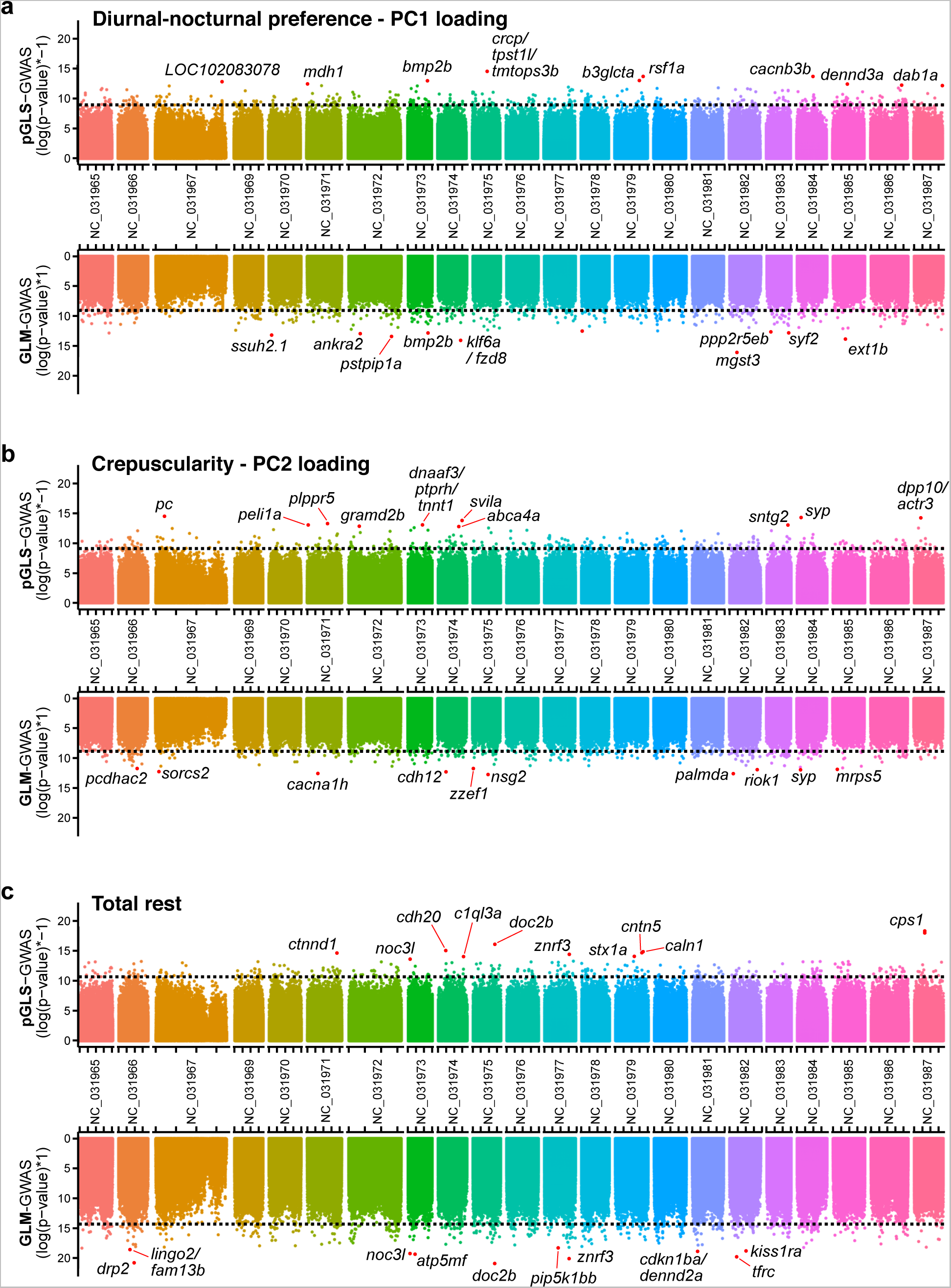
Variants associated with temporal activity patterns are distributed throughout the genome. **a-c,** Manhattan plots showing the p-values for the association test for each SNP for both the pGLS and GLM tests for PC1 loadings (**a**), PC2 loadings (**b**), and total rest (**c**). Dotted lines indicated genome wide cutoffs for identification of HAVs. The top 10 HAV are annotated with the nearby gene(s).

**Extended data Figure 8.**
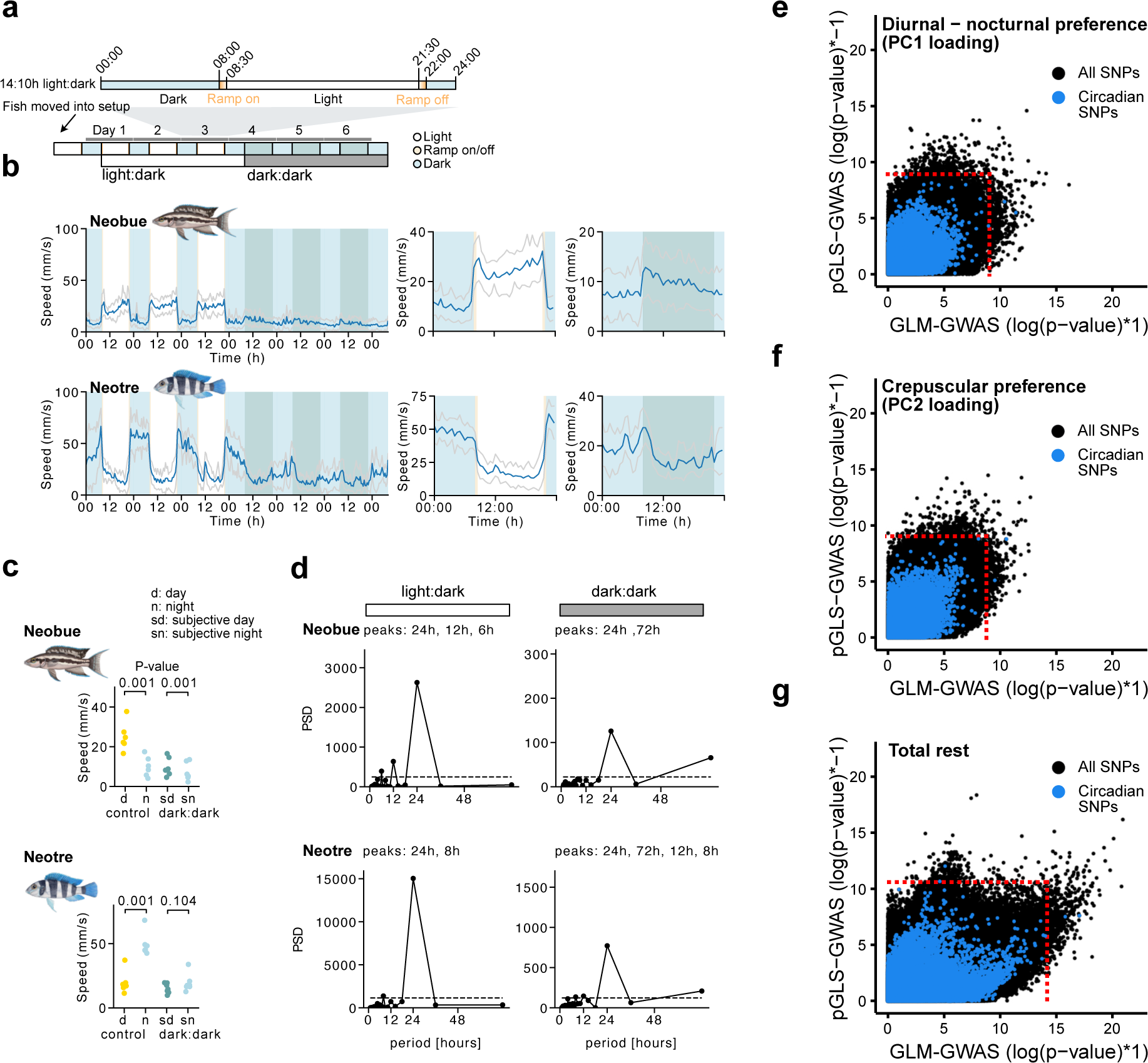
Daily activity patterns are regulated by internal circadian signalling and light. **a,** Schematic of the timeline and light cycle for the behavioural assays of *Neolamprologus buescheri* (Neobue) and *N. tretocephalus* (Neotre). Note that there is a 14:10h light:dark cycle with 3 days of light:dark, and 3 days of dark:dark. **b,** left: weekly speed traces (mean +/- SD) of the full 6 days of assay, right: daily average traces (mean +/- SD) for the light:dark and dark:dark periods. **c,** quantification of speed during the day and night for light:dark, and subjective day and subjective night for the dark:dark period, P-values were calculated by a two-sided paired t-test. **d,** periodograms of the light:dark and dark:dark periods for both species. PSD = power spectral density. Significant peaks are listed on top of the plot for each condition. **e-g**, Scatter plots of the pGLS-GWAS p-value and GLM-GWAS p-values for all SNPs associated with PC1 loadings (e), PC2 loadings (f), and total rest (g). All SNPs associated with circadian genes are labelled. Dotted lines indicated genome wide cutoffs for identification of HAVs.

